# Motor Response Influences Perceptual Awareness Judgements

**DOI:** 10.1101/283762

**Authors:** Marta Siedlecka, Justyna Hobot, Zuzanna Skóra, Borysław Paulewicz, Bert Timmermans, Michał Wierzchoń

## Abstract

Perception and action are closely related, but what is the relation between perceptual awareness and action? In this study we tested the hypothesis that motor response influences perceptual awareness judgements. We designed a procedure in which participants were asked to decide whether a Gabor grating was oriented towards the left or the right. Presentation of the stimuli was immediately followed by a cue requiring a motor response that was irrelevant to the task but could be the same, opposite, or neutral to the correct response to the Gabor patch. After responding to the cue, participants were asked to rate their stimulus awareness using the Perceptual Awareness Scale, and then to report their discrimination decision. Participants reported a higher level of stimulus awareness after carrying out responses that were either congruent or incongruent with a response required by a stimulus, compared to the neutral condition. The results suggest that directional motor response (congruent or incongruent with correct response to the stimulus) provides information about the outcome of decision process and increases the reported awareness of stimuli.

---

Perception and action are closely related, but what is the relation between perceptual awareness and action? The idea that sensorimotor processes might be important for perceptual awareness is not new: it describes awareness as a result of learning sensory (O’Regan & Noë, 2001) and neural (Cleeremans, 2011; Timmermans, Schilbach, Pasquali & Cleeremans, 2012) consequences of actions. Sensorimotor theories of awareness and the enactive approach to consciousness claim that interaction between an organism and its environment shapes its awareness (Noë, 2001; O’Regan & Noë, 2001), but the link between awareness and motor response on the psychophysical level has rarely been investigated.

Other influential theories of consciousness explain the effect of stimulus awareness on stimulus-related behaviour, but do not explicitly expect an influence in the opposite direction. In most of these theories, stimulus awareness depends on the strength of sensory evidence and post-perceptual processing, but the latter is not overtly assumed to be related to the current activity of the motor system. For example, global availability theories claim that a person is aware of a stimulus only if it is represented in a “global workspace” (Baars, 1997; Dehaene & Naccache, 2001; Sergent & Dehaene, 2004). Enough stimuli-related evidence has to be accumulated to cross the threshold of the global availability of information, but the strength of the signal can be additionally affected by attentional processes (Dehaene, Changeux, Naccache, Sackur, & Sergent, 2006). Hierarchical views claim that a person becomes aware of a stimulus when it is represented by a higher-order representation that represents oneself as being in a given first-order mental state (Lau & Rosenthal, 2011; Rosenthal, 2009). This theory allows conscious experience of perceptual stimuli to be based on information other than sensory evidence but does not explicitly predict the influence of ongoing motor activity.

Perceptual awareness can be measured by a number of subjective scales relating to visibility (“continuous scale”, Sergent & Dehaene, 2004) and perceptual awareness (Perceptual Awareness Scale, Ramsøy & Overgaard, 2004), but also by scales that measure perceptual confidence (confidence in one’s perceptual decision, e.g. Cheesman & Merikle, 1986). Rating one’s own awareness of a stimulus is therefore often conceptualized as a decisional process, and researchers aim to describe what information is taken into account during this process. A dominant view is that the judgment of perceptual awareness is determined by stimulus-related information (e.g. Barthelme & Mamassian, 2010; Kiani & Shadlen, 2009; Vickers, 1979). Therefore, researchers focus on studying the characteristics of external stimuli, such as their strength or the type of evidence they provide. It has been suggested that although perceptual decisions are affected by the relative difference between the evidence for each of available responses, confidence in this decision is sensitive mainly to sensory evidence that supports the selected choice, or to absolute evidence for signal over noise (Koizumi, Maniscalco, & Lau, 2015; Samaha, Barrett, Sheldon, LaRocque, & Postle, 2016; Samaha, Iemi, & Postle, 2017; Zylberberg, Barttfeld, & Sigman, 2012).

However, there is some data suggesting that confidence in perceptual decisions might be formed at a late stage of the decision-making process and is based on evidence not available at the time of the stimulus-related decision (Fleming, Maniscalco, Ko, Amendi, Ro, & Lau, 2015; Graziano, Parra & Sigman, 2015). Wierzchoń and colleagues (Wierzchoń, Paulewicz, Asanowicz, Timmermans & Cleeremans, 2014) tested the hypothesis that completing a stimulus-related task influences metacognitive awareness when measured as the relation between task accuracy and awareness ratings. In the experiment, participants were asked to rate stimulus visibility or perceptual confidence either before or after responding to a gender discrimination task. The results showed that both types of awareness measurement predicted discrimination accuracy better when they followed the discrimination response (Wierzchoń et al., 2014). Kiani and colleagues (Kiani, Corthell, & Shadlenet, 2014) showed that level of confidence was related to the time participants took to make the preceding perceptual decision, even though the stimulus strength was kept constant. Other research showed that confidence is sensitive to the outcome of performance monitoring. In an experiment in which participants were asked to decide which of two boxes contained more dots, the level of confidence varied gradually with the magnitude of error-related electrophysiological activity following incorrect response to decisional task (Boldt & Yeung, 2015). This and other studies show that participants can not only differentiate between correct and erroneous responses, but also report confidence that their response was incorrect (Boldt & Yeung, 2015; Charles, Opstal, Marti, & Dehaene, 2013; Scheffers & Coles, 2000). This phenomenon cannot be easily accounted for by theories explaining confidence purely in terms of the accumulation of stimulus-related evidence. Fleming and colleagues (Fleming et al., 2015) provided direct support for the view that the motor system contributes to judgments of perceptual confidence. In their experiment, participants were asked to discriminate between the locations or orientation of two stimuli using their left or right hand and to rate perceptual confidence. Additionally, unilateral single-pulse transcranial magnetic stimulation (TMS) was applied to the dorsal premotor cortex associated with either a chosen or not chosen response, either before or immediately after providing the discrimination response.

The results showed that confidence was influenced by changes in neural activity related to motor response and was lower when the stimulation was incongruent with participants’ correct responses. The effect was similar no matter whether TMS stimulation occurred before or after the discrimination response. In another study, in which electromyographic measure of motor preparatory activity was collected in a perceptual discrimination task, participants reported higher perceptual confidence in trials when sub-threshold motor activation was present before an overt response, even when this activation was not associated with a correct response (Gajdos, Fleming, Saez Garcia, Weindel, & Davranche, 2018).

The idea that motor system activity may contribute to visual confidence is supported by data from neurophysiological studies on perceptual decisions. It has been suggested that the motor system is an integral component of perceptual decision-making processes; in tasks in which stimuli characteristics are directly related to specific, predictable motor reactions, sensory evidence is accumulated directly into a motor response (e.g. Gold & Shadlen, 2003; Hernández, Zainos, & Romo, 2002; Heekeren, Marrett, Bandettini, & Ungerleider, 2003; Shadlen & Newsome, 1996; Spivey, Grosjean, Knoblich, 2005; Wyss, König, & Verschure, 2004). This also happens without conscious perception, i.e. unseen stimuli evoke activation that can be detected at the motor level (Dehaene, 1998; Vorberg, Mattler, Heinecke, Schmidt, & Schwarzbach, 2003). In such cases, motor response itself could provide additional information about one’s own decisional process (Fleming & Daw, 2016), the ease of choice (Kiani et al., 2014), or the outcome of performance monitoring (Boldt & Yeung, 2015).

In the experiment presented in this paper we aimed to test whether motor response influences the report of perceptual awareness of the preceding stimuli. In all the aforementioned experiments concerning response contribution to perceptual awareness, participants were asked to report their confidence in their decisions. However, confidence in one’s own decision could be more sensitive to decision-related and response-related characteristics than perceptual awareness ratings. We used the Perceptual Awareness Scale (PAS) to avoid confusing perceptual awareness with confidence in one’s choice. We also aimed to separate the motor response following stimulus presentation from the stimulus-related decision. In most decisional tasks, motor response, used as an indicator of the decision, is itself indistinguishable from the results of the decision process. Therefore we created a condition in which motor response would be as little “contaminated” by decisional outcomes as possible. To do so we cued a response that was irrelevant to the stimulus-related decision but immediately followed stimulus presentation and directly preceded PAS. This cued response shared the response code with stimulus-related responses. Specifically, we used a discrimination task in which participants were asked to determine whether the Gabor grating was oriented towards the left or the right. Immediately after the Gabor presentation, a cue was presented that required a motor response that was irrelevant to the task but could be the same, opposite, or neutral to the correct response to the Gabor patch. After responding to the cue, participants were asked to rate perceptual awareness of the stimuli (using PAS) and then to report their discrimination decision. Therefore, we created conditions in which cued motor response was either stimulus-congruent, stimulus-incongruent, or neutral. We hypothesized that the cued response would not affect the accuracy of Gabor discrimination, but it would influence the reported awareness of the stimuli. We expected to observe one of three ways in which this response could contribute to the report on perceptual awareness. Firstly, motor response congruent with stimulus-related response could provide additional positive evidence that results in higher perceptual awareness ratings in the Stimulus-congruent condition compared to the other conditions (as observed in the case of the stimulus-related positive evidence effect on confidence, e.g. Zylberberg, Barttfeld, & Sigman, 2012). Secondly, motor activity incongruent with a stimulus-related response could reduce perceptual awareness of the stimulus in the Stimuli-incongruent condition, similarly to the results obtained in a TMS study (Fleming et al, 2015), which would support the hypothesis that disrupting the stimulus-related motor process increases uncertainty about the results of one’s perceptual processing. Lastly, any motor response that potentially overlaps with a stimulus-required response (congruent or incongruent, with correct response to the stimulus) could be interpreted as providing additional information about the decision process and its outcome, thus leading to higher awareness ratings compared to the Neutral condition (similarly to the data showing that any preparatory motor activity increases perceptual confidence, Gajdos et al., 2018).

## Methods

### Participants

Twenty-four healthy volunteers (5 males), aged 21.63 (SD = 2.37) took part in the experiment in return for a small payment. All participants had normal or corrected-to-normal vision and gave written consent to participation in the study. The ethical committee of the Institute of Psychology, Jagiellonian University approved the experimental protocol.

We decided prior to data collection to test at least 20 participants and this sample size was based on previous studies on contribution of motor and pre-motor activation for perceptual confidence (Fleming et al., 2015; Gajdos et al., 2018). No statistical analyses were performed before the completion of data collection.

### Materials

The experiment was run on PC computers using PsychoPy software (Peirce, 2007). We used LCD monitors (1280 x 800 resolution, 60Hz refresh rate). The stimuli were Gabor gratings embodied in visual noise and oriented towards the left or right (45 degrees), presented in the centre of the screen against a grey background. The visual angle of the stimuli was about 3°. The contrast of the stimuli was determined for each participant during a calibration session.

The PAS was presented with the question, “How clear was your experience of the stimulus?”; the options were ‘no experience’, ‘a brief glimpse’, ‘an almost clear experience’, and ‘a clear experience’. The meaning of the individual scale points was explained in the instruction. The description of each point was based on a guide by Sandberg & Overgaard (2015), with some modifications related to the characteristics of the stimuli that were relevant in this experiment (i.e. ‘no experience’ was associated with no experience of the Gabor stripes, but ‘a brief glimpse’ was associated with an experience of ‘something being there’
but without the ability to determine the orientation of the stripes).

### Procedure

The experiment was run in a computer laboratory for four consecutive days in one-hour sessions. All trials began with a blank presentation (500 ms) followed by a fixation cross (500 ms). The grating embedded in white noise was presented for 33 ms. Participants were asked to state whether the grating was oriented towards the left or the right side (using keys “L” and “R” with their left hand).

On the first day, participants started by completing 15 training trials with feedback to get familiar with the stimuli (here presented in colour in RGB space = [0.3,0.3,0.3] and opacity = 1). Then the staircase procedure was used to estimate the stimulus contrast that would lead to about 79% of correct discrimination responses. There were 200 trials with a 1- up 3-down staircase (stair size 0.005, limit for 0.02 and 0.08) and the contrast was established based on the last 150 trials. This was followed by 10 trials in which the PAS scale was presented before discrimination response. Participants used their right hand to report the stimulus visibility (keys 1–4).

Each consecutive session started with a 10-trial training session for the main task; this was followed by 300 experimental trials, which gave 900 experimental trials per participant in total. Each trial started with a central fixation point, after which the Gabor grating was presented. Afterwards participants were asked to respond to the motor cue that was presented in the centre of the screen. The cue was either a vertical bar or an arrow pointing left or right. Participants were asked to press “space” when a vertical bar appeared, “L” when an arrow pointing left was presented, and “R” for an arrow pointing right. The response keys overlapped with those used for Gabor responses, but participants were explicitly told that this task was irrelevant to the main task and were asked to react as quickly and accurately as possible. After participants responded to a cue, the PAS appeared, followed by a discrimination task. The time limit for all responses was 3 seconds. The outline of the procedure is presented in Figure 1.

**Figure 1.**
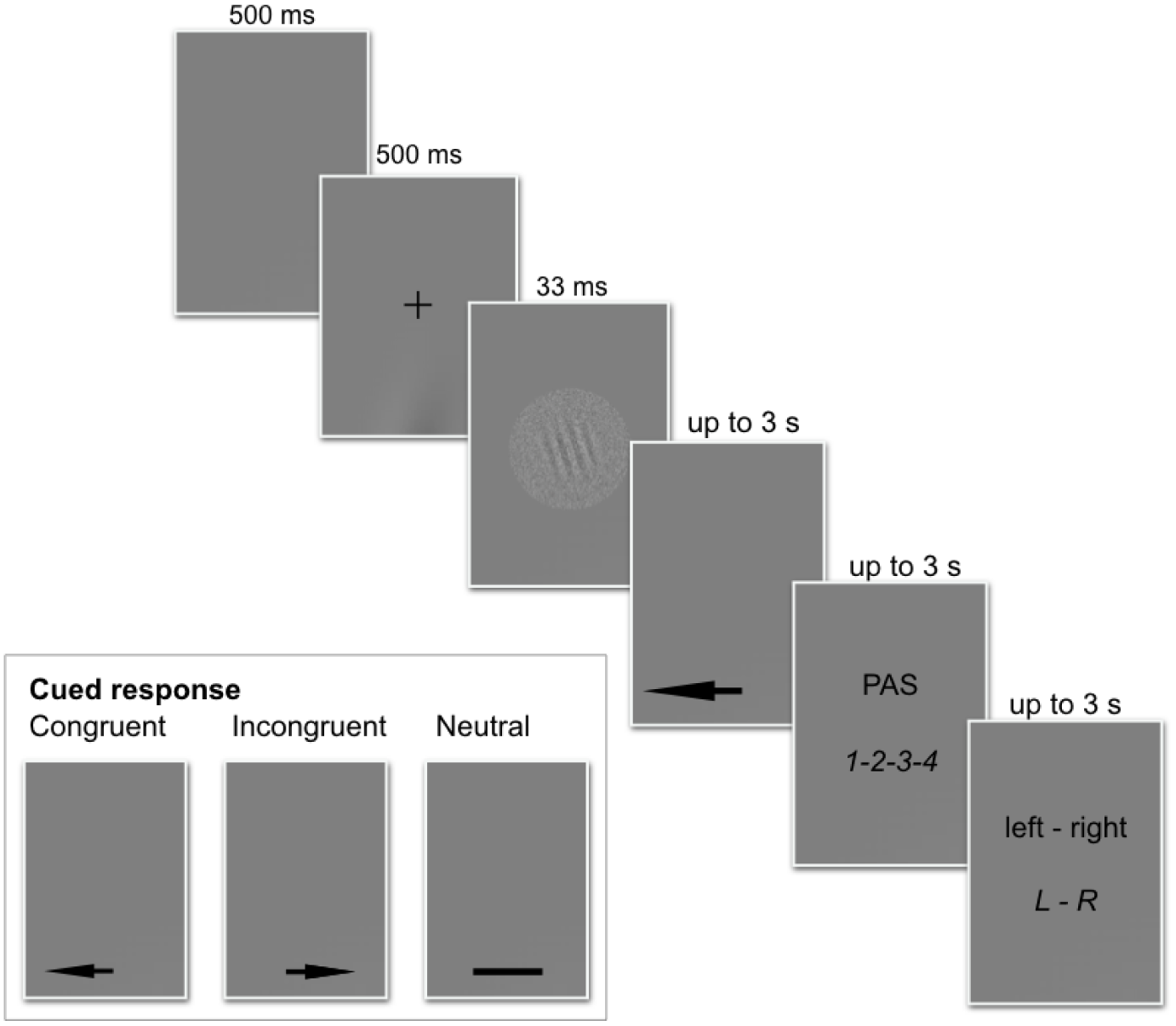
The outline of the experimental procedure. Please note that congruency of the cued response refers to the stimulus in the current trial.

After each session, participants’ accuracy in the motor cue and Gabor discrimination task was estimated, so participants with low accuracy could practise again and be motivated to perform better.

### Results

The main three conditions in the experiment were created by the congruency between the motor cue and correct response to the stimulus: Stimulus-congruent, Stimulus-incongruent, and Neutral (later referred to as Congruent, Incongruent, and Neutral, respectively). We found no statistically significant difference in discrimination accuracy between the conditions (accuracy 78% in all conditions). Also, participants followed the motor cues with a similar efficiency in all conditions (cue-related response accuracy: Congruent – 95%, Incongruent – 94%, Neutral – 95%). Since we were interested only in the trials in which participants followed the motor cue, prior to analysis all trials with incorrect responses to the cue were removed (1,090 trials). In the remaining trials, no significant differences between the conditions were found in respect to the stimuli discrimination accuracy (Congruent – 78%, Incongruent – 79%, Neutral – 78%, *p* > .8). Also signal-detection analysis of responses in the orientation task did not reveal significant differences between conditions in respect to d’ (*p* > .71) or response bias (*p* > .7).

### Confirmatory analyses

To estimate the influence of the cued motor response and discrimination accuracy on the PAS ratings, we used a linear mixed model with random intercept, discrimination accuracy and condition effect (Table 1). The first row (intercept) refers to the average PAS ratings in the baseline condition (Neutral condition, incorrect responses). The second and the third row show that PAS ratings are significantly different in Congruent and Incongruent conditions compared to Neutral condition (for incorrect responses). The forth row estimates the difference in the average PAS ratings between correct and incorrect responses in Neutral condition. The fifth and the sixth rows estimate the interactive effects between accuracy and condition.

**Table 1.** Linear mixed model estimating the effect of condition and accuracy on PAS ratings

| N = 20160 | Estimate | Std. error | df | *t* | *p* | 95% CI |
| --- | --- | --- | --- | --- | --- | --- |
| Intercept | 1.14 | 0.12 | 22.74 | 9.60 | <.001*** | [0.91, 1.38] |
| Stimulus-congruent | 0.17 | 0.36 | 23.43 | 4.66 | <.001*** | [0.10, 0.24] |
| Stimulus-incongruent | 0.13 | 0.04 | 22.59 | 3.53 | .002** | [0.06, 0.20] |
| Accuracy | 0.43 | 0.05 | 22.69 | 8.36 | <.001*** | [0.33, 0.53] |
| Accuracy:Stimulus-congruent | -0.10 | 0.03 | 32.02 | -3.03 | .005** | [-0.17, -0.04] |
| Accuracy:Stimulus-incongruent | -0.03 | 0.04 | 27.09 | -0.84 | .41 | [-0.10, 0.04] |

Likelihood ratio: $\chi^2(25) = 1477, p < .001^*$ $p < .05$ ; \*\* $p < .01$ ; \*\*\* $p < .001$

For better readability of main effects and their interactions we present ANOVA table (Table 2). The average PAS ratings were lower for erroneous than for correct responses within each condition. The contrast analysis showed that in the Neutral condition, PAS ratings were lower than in the other conditions for both correct and incorrect responses (Table 3, Figure 2). We found no differences between Congruent and Incongruent conditions in this respect. There was a significant interaction between Congruency and Accuracy, with one significant contrast: the difference in PAS between the Incongruent and Neutral conditions was smaller for correct responses (*t*(74) = 2.5, *p* = .01).

**Figure 2.**
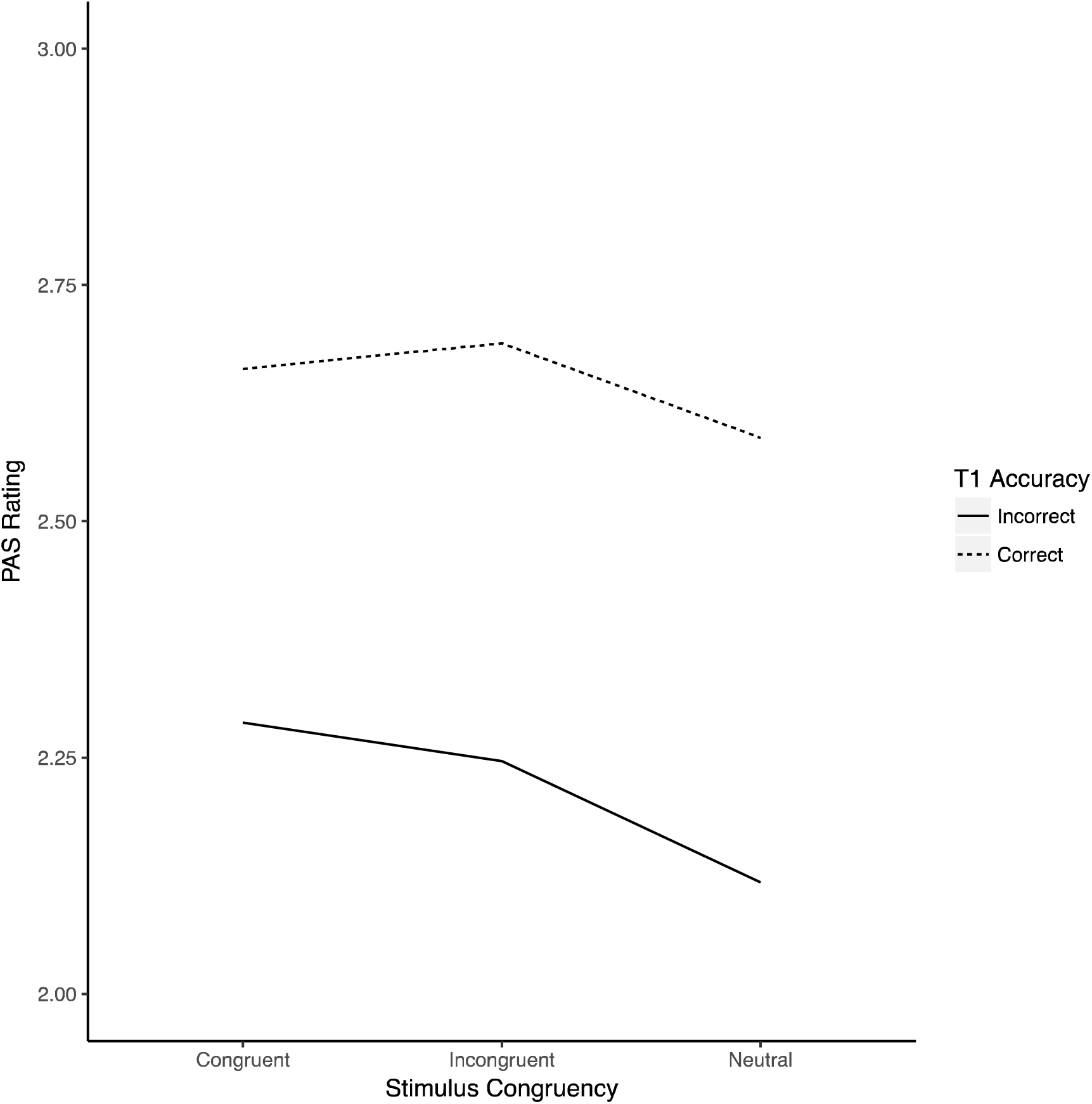
PAS ratings predicted from Stimulus congruency and Accuracy

**Table 2.**
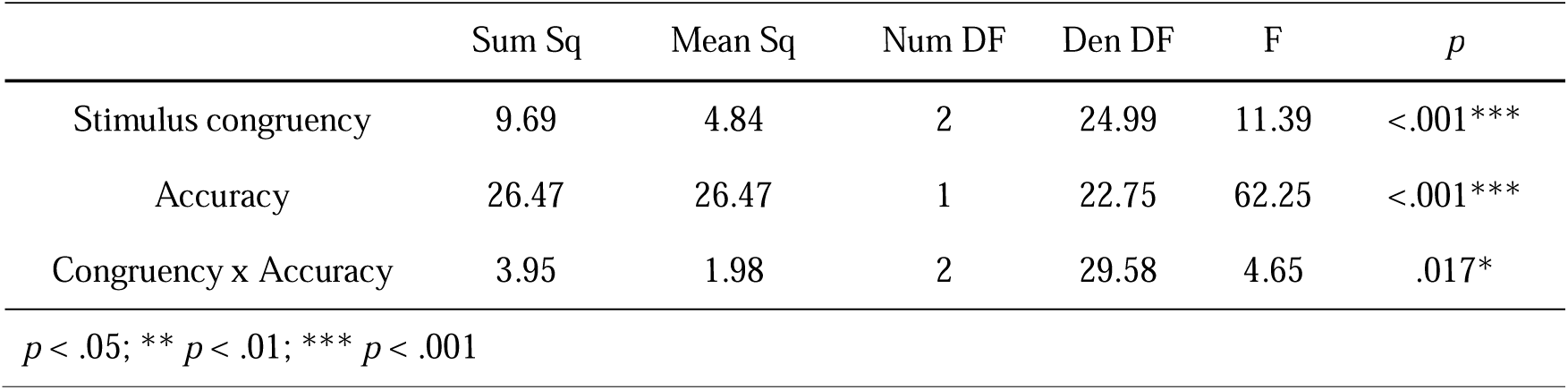
ANOVA table for the linear mixed model predicting PAS ratings from Congruency and Accuracy

|  | Sum Sq | Mean Sq | Num DF | Den DF | F | *p* |
| --- | --- | --- | --- | --- | --- | --- |
| Stimulus congruency | 9.69 | 4.84 | 2 | 24.99 | 11.39 | <.001*** |
| Accuracy | 26.47 | 26.47 | 1 | 22.75 | 62.25 | <.001*** |
| Congruency x Accuracy | 3.95 | 1.98 | 2 | 29.58 | 4.65 | .017* |

$p < .05$ ; \*\* $p < .01$ ; \*\*\* $p < .001$

Table 3

Contrast analyses for the difference in PAS rating level: A. Within conditions, between trials with correct and incorrect discrimination responses; B. Between conditions, separately for correct and incorrect discrimination responses

**A.**
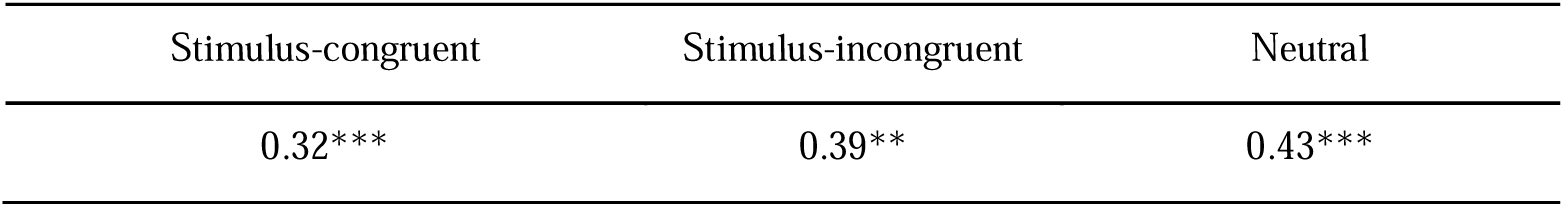
Difference in PAS between correct and incorrect discrimination within conditions

**B.** Difference in PAS between conditions

B. Difference in PAS between conditions
|  | Correct discrimination |  | Incorrect discrimination |  |
| --- | --- | --- | --- | --- |
|  | Stimulus-congruent | Stimulus-incongruent | Stimulus-congruent | Stimulus-incongruent |
| Stimulus-congruent |  | 0.03 |  | -0.04 |
| Neutral | -0.07*** | -0.1*** | -0.17*** | -0.13** |
| *$p < .05$, **$p < .01$, ***$p < .001$ | | | | |

We also compared the frequency of high and low PAS ratings between conditions. All the ratings were encoded as binary outcomes, either high (‘an almost clear experience’ or ‘a clear experience’) or low (‘no experience’ or ‘a vague experience’). Mixed logistic regression analysis revealed that low ratings were given significantly more often in the Neutral condition compared to the other conditions (*z* = 5.9,*p* < .001). The Congruent and Incongruent conditions did not differ between each other (*z* = 1.6, *p* = .1). The frequencies of each PAS rating in all conditions are presented in Figure 3.

**Figure 3.**
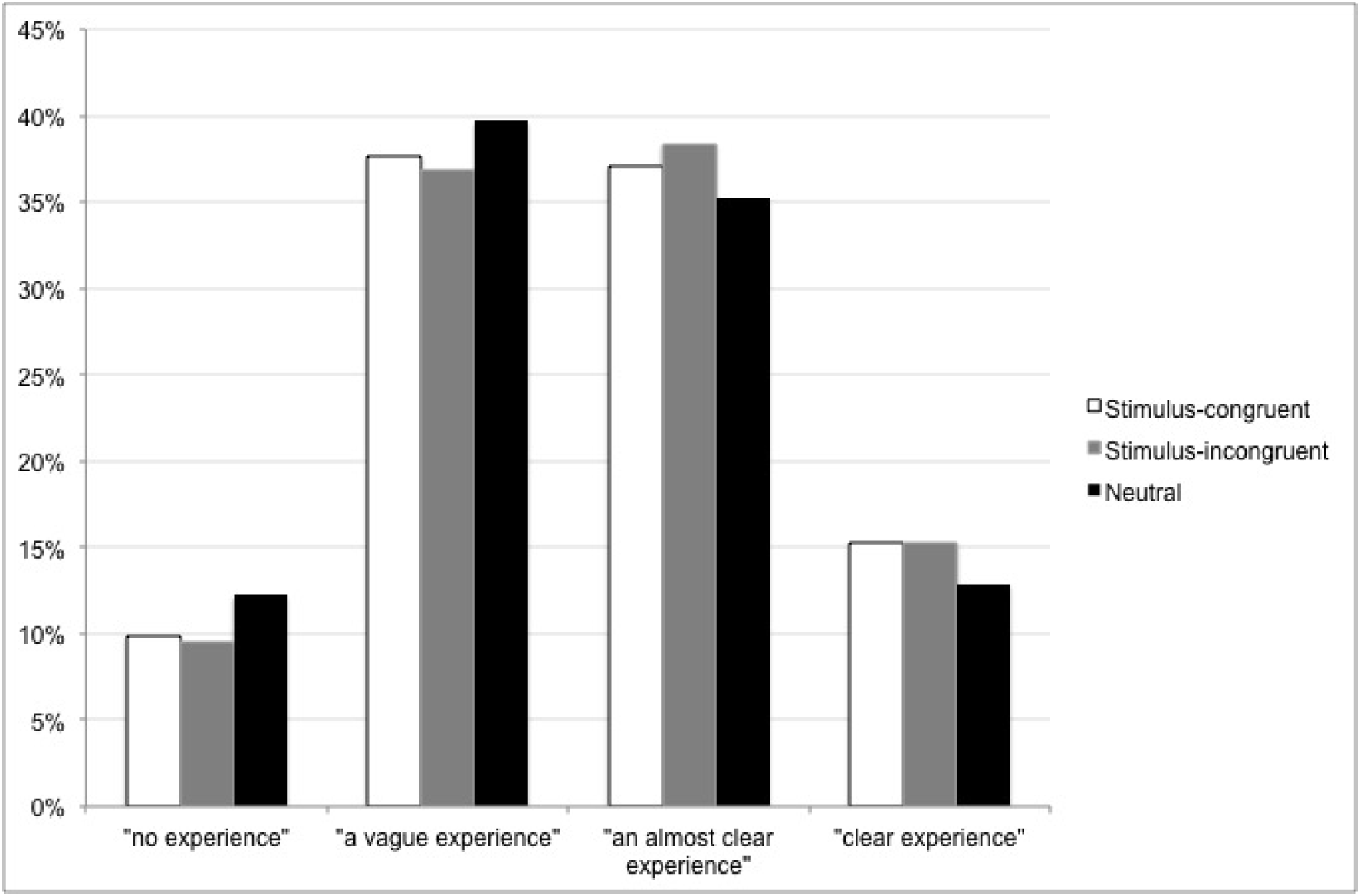
Frequency of each PAS rating in each condition

### Exploratory analyses

Additionally, we analysed reaction times in all three tasks: responding to a motor cue, rating perceptual awareness, and reporting discrimination decision. Participants reacted to the cue fastest in the Neutral condition, and slowest in the Incongruent condition (F(2,46) = 27.90,*p* < .001,*η^2^* = .55; post-hoc Bonferroni analysis showed that all means differed from each other significantly, *p* < .001). In contrast, PAS ratings were given later in the Neutral condition than in the other conditions (F(2,46) = 8.42, *p* = .001, *η_2_* = .27). The post-hoc Bonferroni test showed a significant difference between the Neutral condition and other conditions, *p* = .02, and no significant difference between the Congruent and Incongruent conditions, *p* = .93). Lastly, we found no significant differences between conditions in respect to reaction times in the discrimination task (F(2,46) = 1.25, *p* = .3). Please note that in the above analyses response times were compared only for correct discrimination and cued responses. The average reaction times are presented in Table 5.

**Table 5.** Average reaction times to the cued motor task (ms), PAS rating, and Gabor discrimination task in each condition

| Cued response |  |  | PAS response |  |  | Discrimination response |  |  |
| --- | --- | --- | --- | --- | --- | --- | --- | --- |
| Congruent | Incongruent | Neutral | Congruent | Incongruent | Neutral | Congruent | Incongruent | Neutral |
| *M (SD)* | *M (SD)* | *M (SD)* | *M (SD)* | *M (SD)* | *M (SD)* | *M (SD)* | *M (SD)* | *M (SD)* |
| 1018 (228) | 1065 (243) | 947 (209) | 614 (215) | 608 (218) | 645 (212) | 598 (138) | 587 (115) | 590 (131) |

## Discussion

In this experiment we showed that motor response influences judgments of perceptual awareness of the preceding visual stimulus. Participants were cued to carry out a response immediately after stimulus presentation and – although this response was irrelevant to the main task – it overlapped with the response code to the stimulus discrimination task. The results showed that participants reported a higher level of stimulus awareness after giving a response that was either congruent or incongruent with the response required by the stimulus, compared to the neutral condition.

The results support the hypothesis that a motor response that overlaps with potential responses to the stimuli (i.e. congruent or incongruent with the correct response to the stimulus) provides additional information about the decision process and increases the perceptual awareness of a stimulus. The results are in line with recently published data showing that preparatory activation of thumb muscles is associated with higher confidence in perceptual discrimination, regardless of whether the activation is congruent with a correct response (Gajdos et al., 2018). To our knowledge, no theory of perceptual awareness predicts such an effect. One way of interpreting the results in the context of consciousness theories is in reference to hierarchical approaches which claim that awareness is a result of re-representation of a lower-order state that represents conscious content (Cleeremans 2011; Lau & Rosenthal, 2011; Timmermans et al., 2012). This redescription may be seen as an active process that allows the rebuilding or interpreting of weak representation of conscious content by adding new information, for example from other senses (Łukowska, Sznajder, & Wierzchoń, 2018). In our experiment, motor response following stimulus constitutes additional information that was integrated into perceptual awareness judgment.

An alternative explanation could be proposed that does not assume that action itself influences perceptual awareness. Arrows signalling directional cued response could signal the possibility of increased task difficulty and a conflict between cued response and subsequent discrimination response, compared to the “safe” neutral condition. When following arrow cues, participants could have become more cautious or engaged more deeply in stimuli-related decisional and memory processes that would increase their stimuli recollection or stimuli awareness ratings. Indeed, reaction times to the cue were shortest in the neutral condition, showing it was the easiest; however, the other two conditions also differed in terms of reaction times and we found no significant differences between these conditions in PAS ratings. Also, if after seeing the arrow cue, participants engaged in deep processing of the Gabor patch, we should have observed shorter reaction times to the subsequent orientation decision. Moreover, it seemed that participants were more cautious in the neutral condition when it came to awareness rating: they choose lower scale points more often than in other conditions and their PAS rating latencies were longest compared to other conditions.

In studies on metacognition, a negative relation between the latency of confidence judgment and the level of confidence has been found (e.g. Hilgenstock, Weiss, & Witte, 2014; Pleskac & Busemeyer, 2010; however, “fast guesses” have also been observed, Baranski & Petrusic, 1998; Petrusic & Baranski, 2003). This relation is thought to reflect an additional stage of collecting judgment-related information and indicates the difficulty of reaching the decision. Also, confidence ratings seem to be higher in conditions in which more choice-related information is available, compared to conditions in which it is limited. For example, in a task in which participants solved anagrams and then decided whether a subsequently presented target word was or was not a solution of the anagram, lower confidence ratings and higher frequency of low ratings (cautious strategy) were observed in the condition in which less decision-related information was available, i.e. in the condition in which participants rated their confidence in recognizing the anagram solution before they even saw the target word (Siedlecka, Paulewicz, & Wierzchoń, 2016). It is therefore possible that in our experiment directional cued response, even though it was not directly related to the discrimination task, provided participants with additional information. For example, PAS ratings can be informed by reaction time to arrows together with the experienced ease or difficulty of responding. The analyses of the reaction times to the arrow cues suggest the occurrence of the congruency effect (e.g. Egner, 2017): responses were slower in the incongruent condition compared to the congruent one. This difference suggests that presentation of the Gabor patch automatically activated a motor plan related to the orientation task that either facilitated or was in conflict with the following cued response. Recently, Fleming and Daw (2017) proposed a hierarchical model of metacognition that assumes that confidence judgments are informed by one’s actions. In this model, a second-order level assesses not only the internal sensory evidence for the decision, but also one’s performance (e.g. by detecting errors). Although the model relates explicitly to confidence in one’s decisions, the authors claim that it could apply to different types of self-evaluation.

One interesting area for future exploration is the relation between error monitoring and stimulus awareness processes. It has been suggested that monitoring processes evaluate ongoing performance and correct one’s errors without engaging conscious processing, i.e. even when errors remain unnoticed due to the speed of responses or when participants cannot intentionally monitor their performance due to stimuli degradation (Endrass, Reuter, & Kathmann, 2007; Logan & Crump, 2010; Nieuwenhuis, Ridderinkhof, Blom, Band, & Koket, 2001; Nieuwenhuis, Schweizer, Mars, Botvinick, & Hajcak, 2007; Wessel, Danielmeier, & Ullsperger, 2011). In speeded response tasks, error-related neural activity seems to result from a comparison between the representation of the correct response and the response actually given (Bernstein, Scheffers, & Coles, 1995). The results of performance monitoring could potentially influence perceptual awareness. However, post-error slowing is usually observed after an error is detected by monitoring processes. In our experiment we did not detect any delay in PAS ratings in the incongruent condition.

Summing up, in this experiment we showed that judgments of stimulus awareness could be influenced by a preceding motor response. Future studies are needed to determine whether perceptual awareness judgments are sensitive to lower-order senso-motor processes or whether motor response-related characteristics inform awareness judgment. Nevertheless, this finding represents an empirical hurdle for theories of perceptual awareness.

